# Age-related changes to macrophages are detrimental to fracture healing

**DOI:** 10.1101/720128

**Authors:** Daniel Clark, Sloane Brazina, Frank Yang, Diane Hu, Erene Niemi, Ted Miclau, Mary Nakamura, Ralph Marcucio

## Abstract

The elderly population suffers from higher rates of complications during fracture healing that result in increased morbidity and mortality. Inflammatory dysregulation is associated with increased age and is a contributing factor to the myriad of age-related diseases. Therefore, we investigated age-related changes to an important cellular regulator of inflammation, the macrophage, and the impact on fracture healing outcomes. We demonstrated that old mice (24 months) have delayed fracture healing with significantly less bone and more cartilage compared to young mice (3 months). The quantity of infiltrating macrophages into the fracture callus was similar in old and young mice. However, RNA-seq analysis demonstrated distinct differences in the transcriptomes of macrophages derived from the fracture callus of old and young mice, with an upregulation of M1/pro-inflammatory genes in macrophages from old mice as well as dysregulation of other immune-related genes. Preventing infiltration of the fracture site by macrophages in old mice improved healing outcomes, with significantly more bone in the calluses of treated mice compared to age-matched controls. After preventing infiltration by macrophages, the macrophages within the fracture callus were collected and examined via RNA-seq analysis, and their transcriptome resembled macrophages from young calluses. Taken together, infiltrating macrophages from old mice demonstrate detrimental age-related changes, and depleting infiltrating macrophages can improve fracture healing in old mice.

## 1. Introduction

Fracture healing follows a distinct temporal sequence characterized by an initial inflammatory phase followed by anabolic and catabolic phases [1]. The inflammatory phase is characterized by recruitment of innate and adaptive immune cells to the fracture site [2]. Molecular interactions among immune cells regulate promotion and resolution of inflammation, as well as recruitment of appropriate progenitor cells necessary during the anabolic phases of fracture healing [3, 4]. Age-related changes in the inflammatory response and the effect on fracture healing are not well-studied.

Perturbation of the inflammatory phase of fracture repair can have detrimental effects on the healing outcome [4]. This is evident in patients with chronic inflammatory conditions such as diabetes, rheumatoid arthritis, and increased age. The disturbance of inflammation in these conditions is associated with poorer fracture healing outcomes [5] [6–8]. Similarly, in experimental animal models, induced local and systemic inflammation has negative effects on fracture healing outcomes [9–11].

A higher incidence of bone fractures occurs in the elderly. The residual life time fracture risk in individuals over 60 years of age is 29% in men and 56% in women [12]. Further, bone fractures in the elderly are associated with increased rates of delayed unions and non-unions with a resulting increase in, morbidity and mortality [13–16]. Fracture models using rodents also have shown delayed healing in old mice compared to young [17, 18]. Therefore, understanding how dysregulation of the inflammatory process in elderly populations affects fracture healing represents a critical area for investigation.

The elderly population, including those in good health, are found to have higher levels of circulating pro-inflammatory cytokines, which is associated with a predisposition to a range of systemic disease including osteoporosis, Alzheimer’s disease, Type II diabetes, atherosclerosis, and Parkinson’s disease [19–21]. The chronic, increased pro-inflammatory status associated with aging is described as “inflamm-aging.” Inflamm-aging has been suggested to result from inadequate resolution of inflammation or the result of chronic stimulation that prolongs the inflammatory response [22, 23]. Currently, the mechanisms responsible for inflamm-aging are unclear, but age-related changes to key cellular regulators of inflammation may be responsible. We have observed increased and sustained systemic inflammation in fracture healing in old animals [24]. Further, in previous research from our laboratory, ablation of the hematopoietic stem cells of old animals via lethal irradiation, and subsequent replacement with hematopoietic stem cells of juveniles, accelerated fracture healing compared to the chimeras that received old bone marrow. This effect was associated with reduced systemic inflammation in the heterochronic chimeras [24]. Thus, manipulating the inflammatory system in old animals could affect the rate of fracture healing.

The macrophage is an important inflammatory cell involved in fracture healing [25, 26]. Throughout the course of healing, macrophages polarize among various states of inflammation in response to their environment. During early phases of healing macrophages are classically activated and exhibit pro-inflammatory activities. These have been generally considered M1 macrophages [27, 28]. As healing progresses macrophages switch to anti-inflammatory states and are generally considered M2 macrophages that are responsible for downregulating inflammation and promoting healing [28, 29]. However, these are very broad categories and not strict definitions of cell types, and in actuality these categories are likely comprised of multiple subsets of macrophages comprising these populations. Nonetheless, enhancement of M2-like macrophages at the fracture site has been shown to improve fracture repair [30]. In addition, tissue-resident macrophages, osteomacs, have been observed in close proximity to osteoblasts on the bone surface, and contribute to osteoblast regulatory functions [31]. Macrophages promote osteoblast differentiation during fracture healing [32], and we have also shown the importance of macrophages in fracture healing. Fracture healing in mice that lack *C-C Motif Chemokine Receptor 2* (*Ccr2*) exhibits disruption of macrophage trafficking to the fracture callus and delayed fracture healing [33], and others have observed similar results after depletion of macrophages in mouse models [25, 30, 32]. Finally, age-related changes to macrophage activity have been previously demonstrated. Dysregulated chemokine and cytokine expression has been observed in aged macrophages compared to young [34]. Additionally, decreased growth factor production is associated with aged macrophages [35]. Hence, these changes could significantly impact fracture healing in aged animals.

In this work, our goal was to better understand the contribution of the inflammatory response in aged animals, and specifically the macrophage, during fracture healing. We assessed influx of inflammatory cells to the fracture site in young and elderly mice and used next-generation RNA sequencing to assess age-related changes in the transcriptome of macrophages derived from the fracture callus. Finally, we manipulated macrophages in elderly mice to asses the extent to which they can be targeted for therapy. Our results add to the increasing body of evidence supporting a role for the inflammatory system in bone fracture healing.

## 2. Materials and methods

### 2.1. Animals

All procedures were approved by the UCSF Institutional Animal Care and Use Committee and conducted in accordance with the National Institutes of Health guidelines for humane animal care. All mice (C57B6/J) were obtained from the National Institute on Aging’s Aged Rodent Colony. Specific pathogen-free mice were bred and raised in barriers and utilized for experiments at 24 months (old adult mice) or 3 months (young adult mice) of age.

### 2.2. Tibia fractures

Mice were anesthetized and subjected to closed, non-stable fractures of the right tibia created by three-point bending, as previously described [36]. Analgesics were administered post-surgery and mice were permitted to ambulate freely. To inhibit macrophage recruitment during fracture healing in old mice, treatment groups received the compound PLX3397 (275mg/kg) (Plexxikon Inc. Berkeley, CA) *ad libitum* in their chow. Control groups received the same chow without PLX3397. PLX3397 is a small molecule kinase inhibitor of Colony stimulating factor 1 receptor (M-CSF1R) [37]. The compound significantly reduces the quantity of macrophages in the fracture callus [38]. Treatment with PLX3397 was started 24 hours before fracture and continued for 3 or 10 days after fracture until the animal was euthanized and the tissues harvested for analysis.

### 2.3. Tibia processing and stereology

Mice were sacrificed at day 10-post fracture for stereological analysis of the fracture callus. Fractured tibiae were collected and fixed for 24 hours in 4% paraformaldehyde. The tibiae were decalcified in 19% EDTA for 14 days and dehydrated in graded ethanol prior to paraffin embedding. Serial sagittal sections (10μm) were cut through the entire tibia using a microtome (Lecia, Bannockburn, IL) and mounted on slides. Sections were stained using Hall Brunt Quadruple Stain (HBQ) to visualize bone and cartilage. An Olympus CAST system (Center Valley, PA) and software by Visiopharm (Hørsholm, Denmark) was used to quantify tissue volumes according to stereological methods developed by Howard and Reed [39]. A 2x magnification setting was utilized to outline the boundary of the fracture callus. Then, bone and cartilage were identified and labeled at 20x magnification. The Cavalieri formula was used to estimate the absolute volume of the total callus, bone, and cartilage tissue as previously described [17, 24, 33].

### 2.4. Flow cytometry

Old and young mice were sacrificed at days 1, 3, 10, and 14 post fracture. Fractured tibiae were collected and the calluses were dissected, weighed, disassociated manually through a 100μm nylon cell strainer, and digested with Collagenase type I (0.2mg/mL; Worthington, Lakewood, NJ) for 1 hour at 37 degrees. Cells were rinsed, collected by centrifugation, and resuspended in incubation buffer (0.5% BSA in PBS). Isolated cells were blocked for 10 minutes at room temperature in 10% rat serum and then stained with directly conjugated fluorescent antibodies: CD45 (clone 30-F11), CD3 (145-2C11), B220 (RA3682), Gr-1 (RB6-8C5), NK1.1 (PK136), MHC Class II (M5/114.15.2) Ly6 C (HK1.4) Ly6G (clone 1A8), F4/80 (clone BM8), and CD11b (clone M1/70) (Biolegend, San Diego, CA). Staining with Fixable Red Dead (Thermo Fisher, Waltham, MA) was used for the detection of dead cells. Isotype controls and fluorescence minus one controls were used to gate for background staining. Cells were sorted on a FACSAria (BD Biosciences, San Jose, CA), and FlowJo Software 9.6 (Treestar, Ashland, OR) was used for analysis.

### 2.5. RNA-seq Analysis

Macrophages were isolated from the fracture callus of old (n=10), old treated with PLX3397 (n=6) and young (n=11) mice at day 3 post-fracture. The callus was dissected and cells were collected as described above. For the detection and isolation of macrophages, cells that stained with the following directly conjugated florescent antibodies CD3 (145-2C11), B220 (RA3682), NK1.1 (PK136), Ly6G (clone 1A8) were excluded, and macrophages were collected by staining with CD45 (clone 30-F11), F4/80 (clone BM8), and CD11b (clone M1/70) (Biolegend, San Diego, CA). Callus macrophages were sorted to 99.8% purity on FACSAria (BD Biosciences, San Jose, CA). RNA was extracted using Invitrogen RNA aqueous Micro Kit (AM1931). The library was prepared using Illumina Truseq Stranded mRNA Library Prep Kit and Single-end 50 bp RNAseq was performed on Illumina HiSeq 4000. An average read depth of 60.7 million reads per sample was generated. Reads were aligned using STAR_2.4.2a to the mouse genome (Ensemble Mouse GRCm38.78). Differential gene expression was assessed using DEseq2. Gene ontology and KEGG pathway analysis was performed using DAVID (http://david.abcc.ncifcrf.gov) and the mouse genome was used as background.

### 2.6. Statistics

GraphPad Prism v.7 software was used for analysis. Comparisons between groups was made by first using a 2-way ANOVA multiple comparisons test followed by a 2-tailed Student’s *t* test. P<0.05 was considered statistically significant. Differential gene expression was considered significant at FDR<0.1. For term enrichment in gene ontology and KEGG pathway analysis the level of significance was set using a modified Fisher Exact P-value of p<0.05.

## 3. Results

### 3.1. Old mice demonstrate delayed fracture healing compared to young mice

Previously, we have shown that the rate of fracture healing is directly related to the age of mice. Eighteen month-old mice healed slower than 1 and 6 month old mice [17]. Therefore, to begin this work we compared fracture healing in 24 month-old mice, because these are considered elderly [40], to young adult mice (3 months old). Fracture healing was assessed using stereology to quantify the volume of bone and cartilage within the fracture callus. At 10 days post-fracture, old mice had smaller calluses with significantly less bone and more cartilage (p<0.05) compared to young adult mice (Fig. 1). Thus, these data are in agreement with our earlier work.

**Figure 1:**
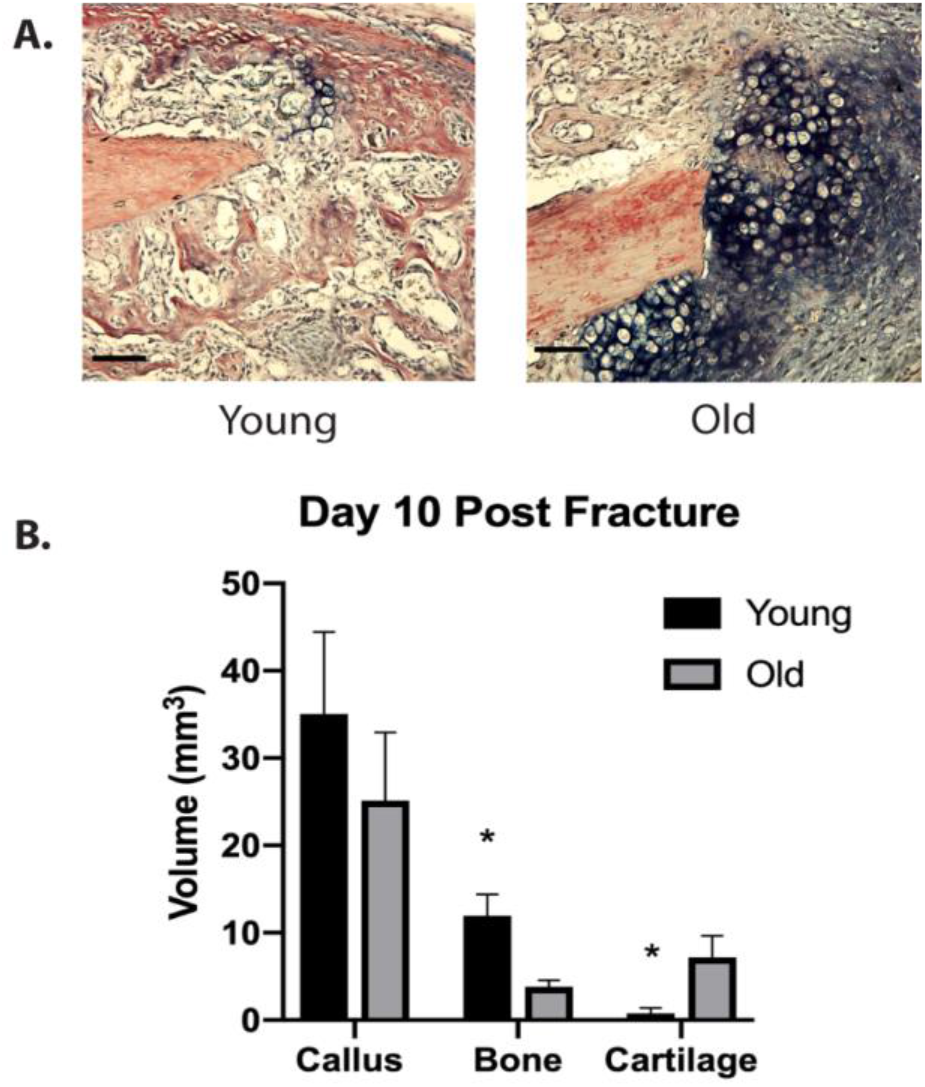
Old mice demonstrate delayed fracture healing compared to young. (A) Representative histological images (HBQ stain) of fracture calluses in old (24 months) and young (3 months) mice (scale bar= 200um). Stereological analysis was performed and the volume of the total callus and the volume of bone and cartilage tissue within the callus was calculated at 10 days after closed tibial fracture (n=5/group). (B) Old mice demonstrate delayed healing with smaller callus size and significantly less bone and more cartilage (* p<0.05).

### 3.2. Immune cell infiltration into the fracture callus is similar in old and young mice

Our objective was to examine the effect of age on inflammation during fracture healing. First, we assessed the inflammatory response during fracture healing in old and young mice by quantifying lymphocyte infiltration into the fracture callus at 1, 3, 10, and 14 days post fracture via flow cytometry. The quantity of T cells, natural killer T cells, and natural killer cells isolated from the callus was similar in young and old mice at all time points examined (n=5 mice/ group) (Fig. 2A). In contrast, B cells were significantly increased in young mice at day 10 (p<0.05). Macrophages were the most prevalent immune cell analyzed among cells derived from the fracture callus. The quantity of macrophages peaked 3 days after fracture, and the macrophages were reduced dramatically by day 14, but, no significant differences were noted in the quantity of F4/80+ macrophages in old and young fracture calluses at any time point analyzed (Fig. 2B). However, in examining subpopulations of macrophages, the F4/80+, Ly6C- population, indicative of a “restorative” macrophage phenotype [41] was increased within the fracture callus of young mice compared to old mice at day 1, but this difference had resolved by day 3 (Fig. 2C). Our data suggest that there may be slight differences in the cellular inflammatory response in young and old mice, but the differences are subtle.

**Figure 2:**
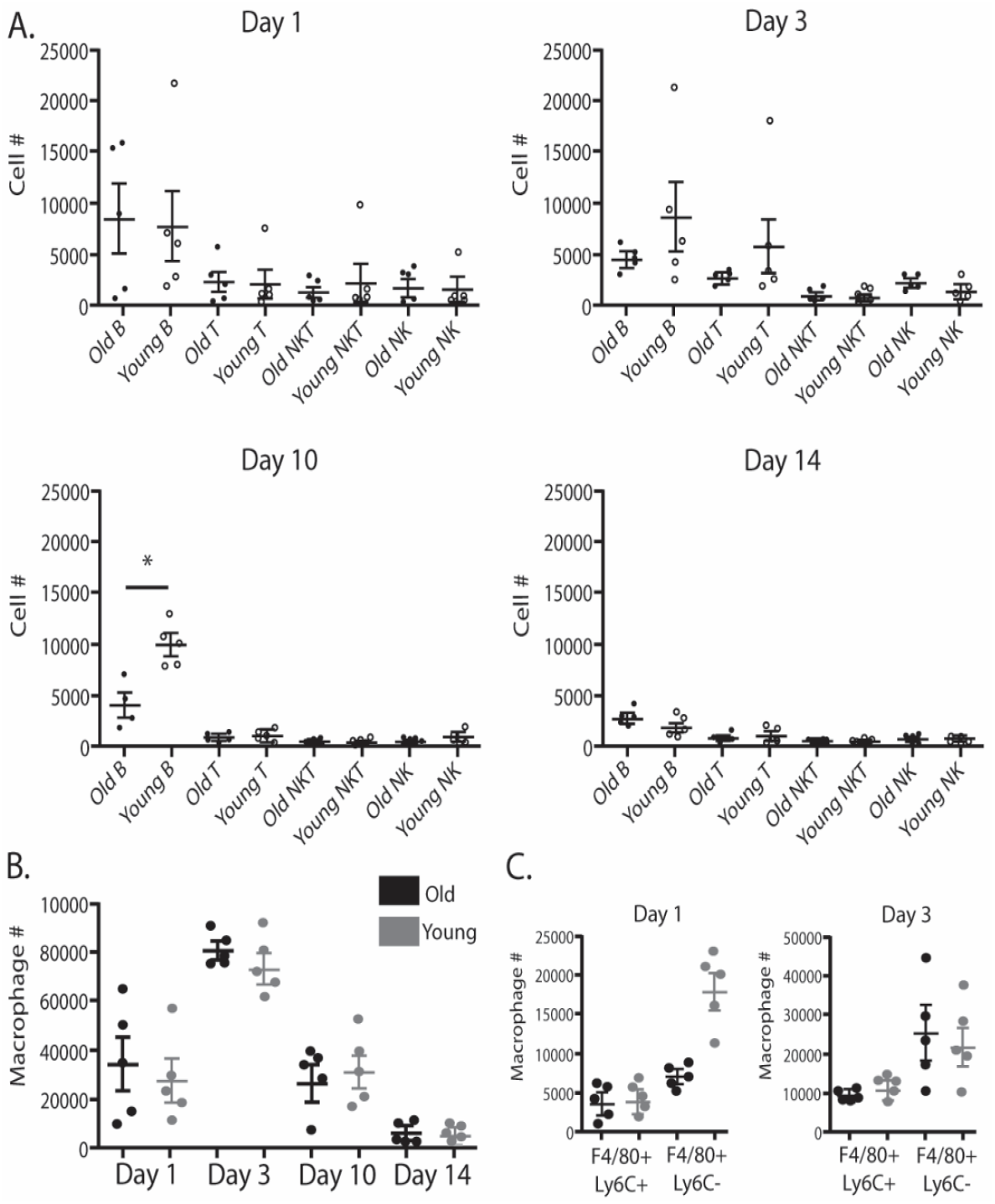
Immune cell infiltration into the fracture callus is similar in old and young mice. (A) The quantity of B cells, T cells, NKT cells, and NK cells were similar within the fracture callus at days 1, 3, 10 and 14 post fracture measured via flow cytometry in old (n=5) and young (n=5) mice. B cell quantity at day 10 was the only significant differences between age groups. (B) Macrophages (F4/80+) were the most abundant immune cell analyzed within the fracture callus and demonstrated no significant difference in quantity between young (gray) and old (black) mice at any of the time points analyzed. (C) A sub-population of macrophages (F4/80+, Ly6C-) was increased in young mice at day 1 compared to old mice. (*p<0.005).

### 3.3. Callus macrophages from old mice are transcriptionally distinct from callus macrophages from young mice

Since the quantity of immune cells infiltrating the fracture callus was similar in young and old mice, functional, rather than quantitative, changes in these cells may contribute to inflammatory dysregulation upon aging. The macrophage was selected for further analysis, because macrophages were the most abundant immune cell type analyzed, and we observed differences in a subpopulation of macrophages, F4/80+,LyC-, between young and old mice. To evaluate intrinsic age-related changes in macrophages, RNA-seq analysis was performed on macrophages isolated from the fracture callus of old and young mice at 3 days post fracture. In total, 1222 genes were significantly differentially expressed in old macrophages compared to young; 364 genes were upregulated and 200 genes were down-regulated more than 2-fold (Fig. 3A). Gene ontology enrichment analysis was performed to begin exploring the implications of these differentially expressed genes in macrophages (Fig. 3B). A number of significantly enriched disease processes are identified that are related to aging and the immune response. These enriched terms include rheumatoid arthritis, graft versus host disease, inflammatory bowel disease. Additionally, molecular and cellular processes important in macrophage function were significantly enriched, including antigen processing and presentation, response to wounding, and cytokine activity. These suggest that fundamental differences in function of young and old macrophages may contribute to age-related differences in inflammatory response to fracture.

**Figure 3:**
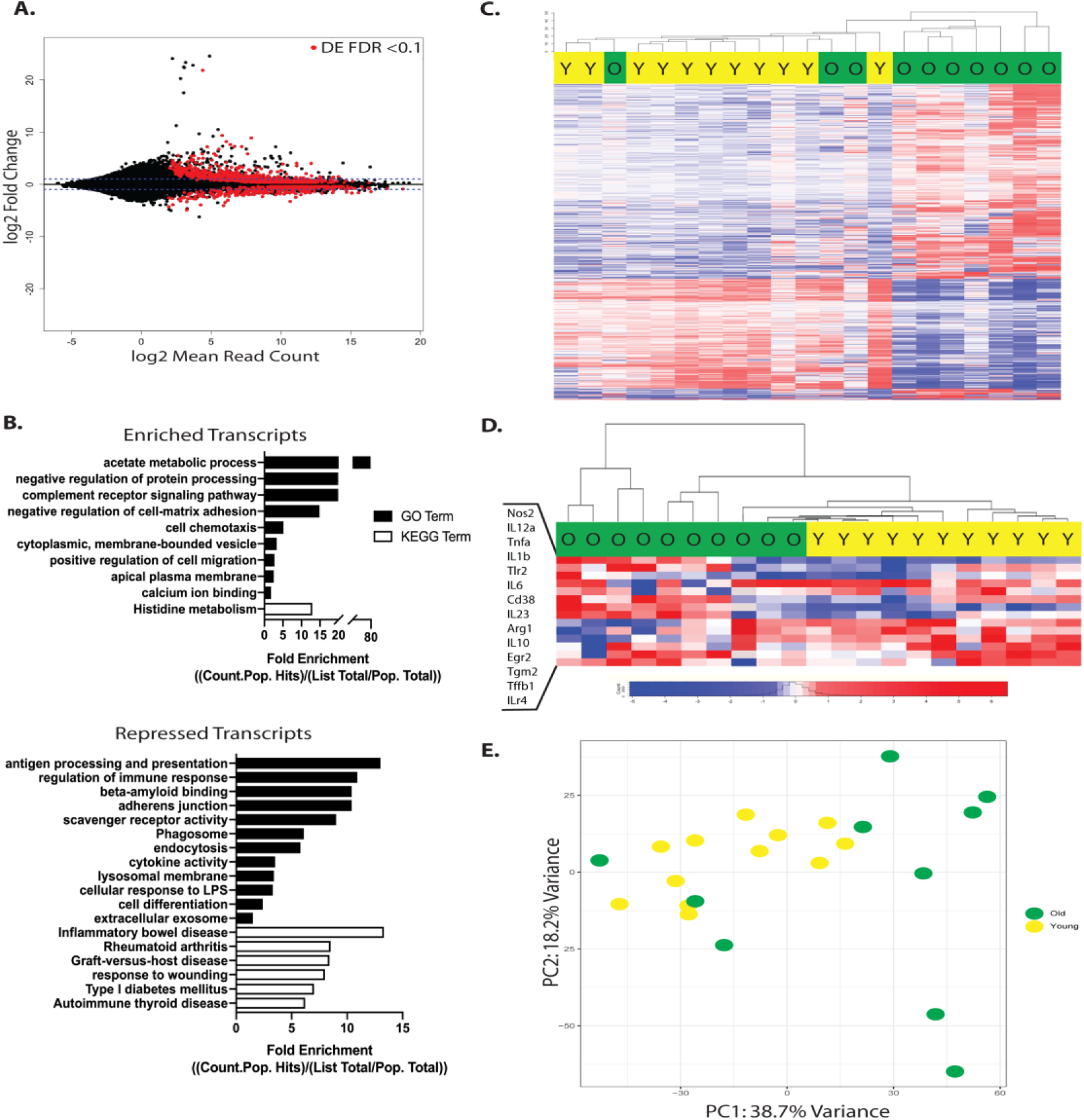
Macrophages from the fracture callus of old mice are transcriptionally distinct from young mice. RNA-seq analysis of callus macrophages in old (n=10) and young (n=11) mice collected at 3 days post fracture. (A) 1222 genes were significantly differentially expressed in old macrophages compared to young (FDR <0.1) (red dots). (B) Enriched gene ontology terms associated with the significantly up and down regulated genes. (C) Heat map demonstrates hierarchical clustering of young (yellow) and old (green) mice based on the differential expression of the 1222 genes. (D) Hierarchical clustering of young (yellow) and old (green) mice, based on the differential expression of a M1/M2 gene signature, demonstrates a more pro-inflammatory/M1 gene expression signature in macrophages from old mice compared to young. (E) Principal component analysis of differential gene expression in macrophages form old mice and young demonstrates clustering of young (yellow) mice and a heterogenous spread of old (green) mice across PC1 and PC2.

The 1222 significantly differentially expressed genes in the macrophages from old and young fracture calluses is compared with a heat map (Fig. 3C). Hierarchical clustering based on Euclidean distances demonstrates old and young animals sort largely based upon differential gene expression patterns (Fig. 3C). The heatmap further characterizes old mice as more heterogeneous in their transcriptomes than young animals. To further assess differences between old and young macrophages, samples were hierarchically sorted based on their differential expression of 14 genes associated with characteristic macrophage cytokines and markers of M1 and M2 macrophages. This analysis demonstrated that mice sort by age and that old mice have increased expression of pro-inflammatory cytokines and markers of M1 macrophages (Fig. 3D). Principal component analysis further demonstrates distinct clustering of the young mice from old (Fig. 3E). The heterogeneity of the old macrophage transcriptome is further seen in the principal component analysis with young mice demonstrating closer clustering compared to old mice that span PC1 and PC2 (Fig. 3E). Median PC scores of old and young mice across the individual principal components additionally demonstrate the difference by age and the increased variability in the old mice (Supplementary Fig. 1). PC1 appears to largely separate the old from young mice. Although, three old mice are seen clustering with the young mice in all analyses performed, further supporting the idea that cells from old mice are more variable in their gene expression profiles than young mice. Genes with the highest and lowest eigenvalues on PC1 are presented in Supplementary Table 1.

### 3.4. Inhibition of macrophage recruitment improves fracture healing in old mice

Macrophages from the fracture calluses of old mice were transcriptionally distinct and displayed a more pro-inflammatory phenotype compared to young macrophages. Thus, we sought to inhibit macrophage recruitment during fracture healing to assess if healing outcomes could be improved in old mice. We administered a CSF-1R inhibitor, PLX3397, that inhibits recruitment of macrophages from the bone marrow. Administering PLX3397 for 10 days after fracture healing improved fracture healing outcomes (Fig. 4). Stereological analysis demonstrated a larger fracture callus with significantly increased bone volume in treated old mice compared to control old mice (Fig 4B). Flow cytometry demonstrated significant reduction of macrophages within the callus of PLX3397 treated mice (Fig. 5A).

**Figure 4:**
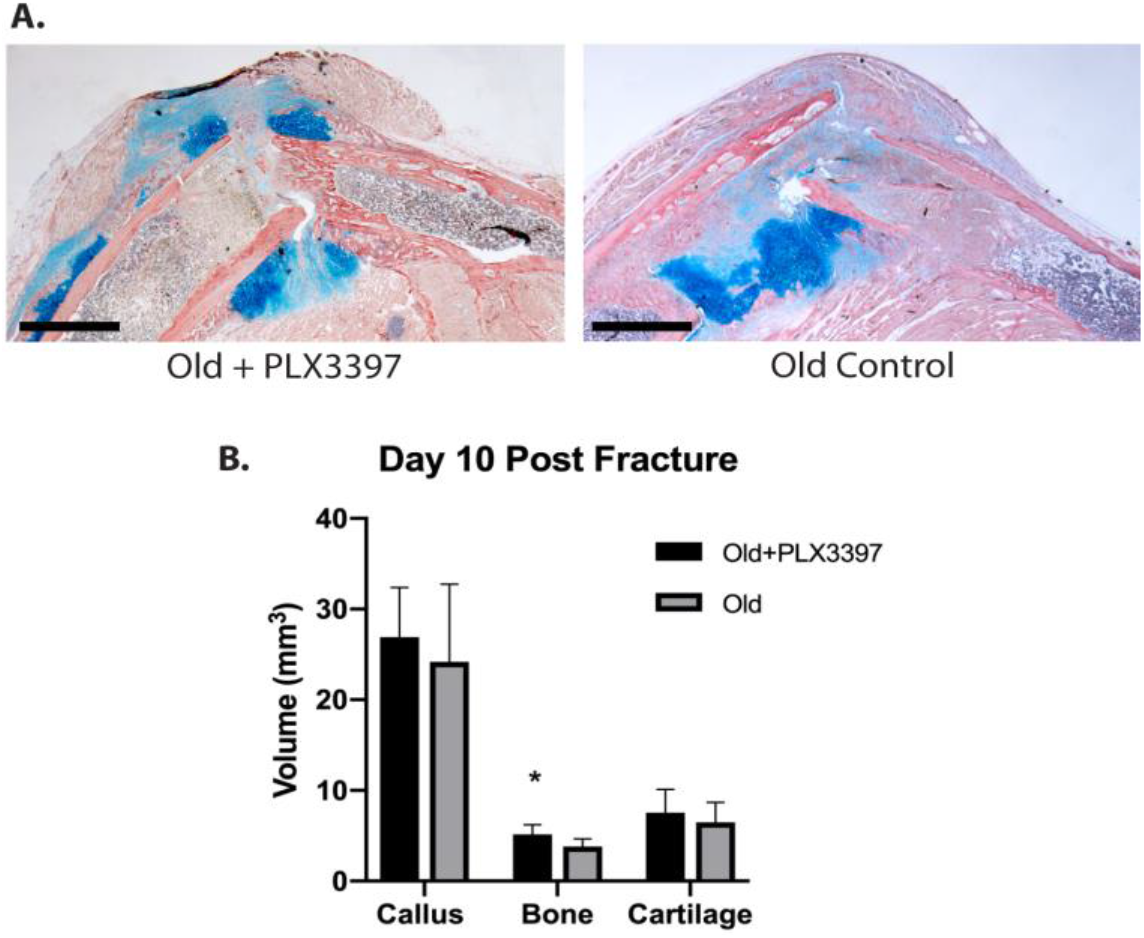
Inhibition of macrophage recruitment improves fracture healing in old mice. (A) Representative histological images (HBQ stain) of fracture calluses in old mice treated with PLX3397 and aged matched old controls 10 days after closed tibial fracture (n=11/group) (scale bar= 200um). Stereological analysis was performed and the volume of the total callus and the volume of bone and cartilage tissue within the callus was calculated. (B) Old mice treated with PLX3397 demonstrate significantly more bone volume compared to old non-treated mice (* p<0.05).

**Figure 5:**
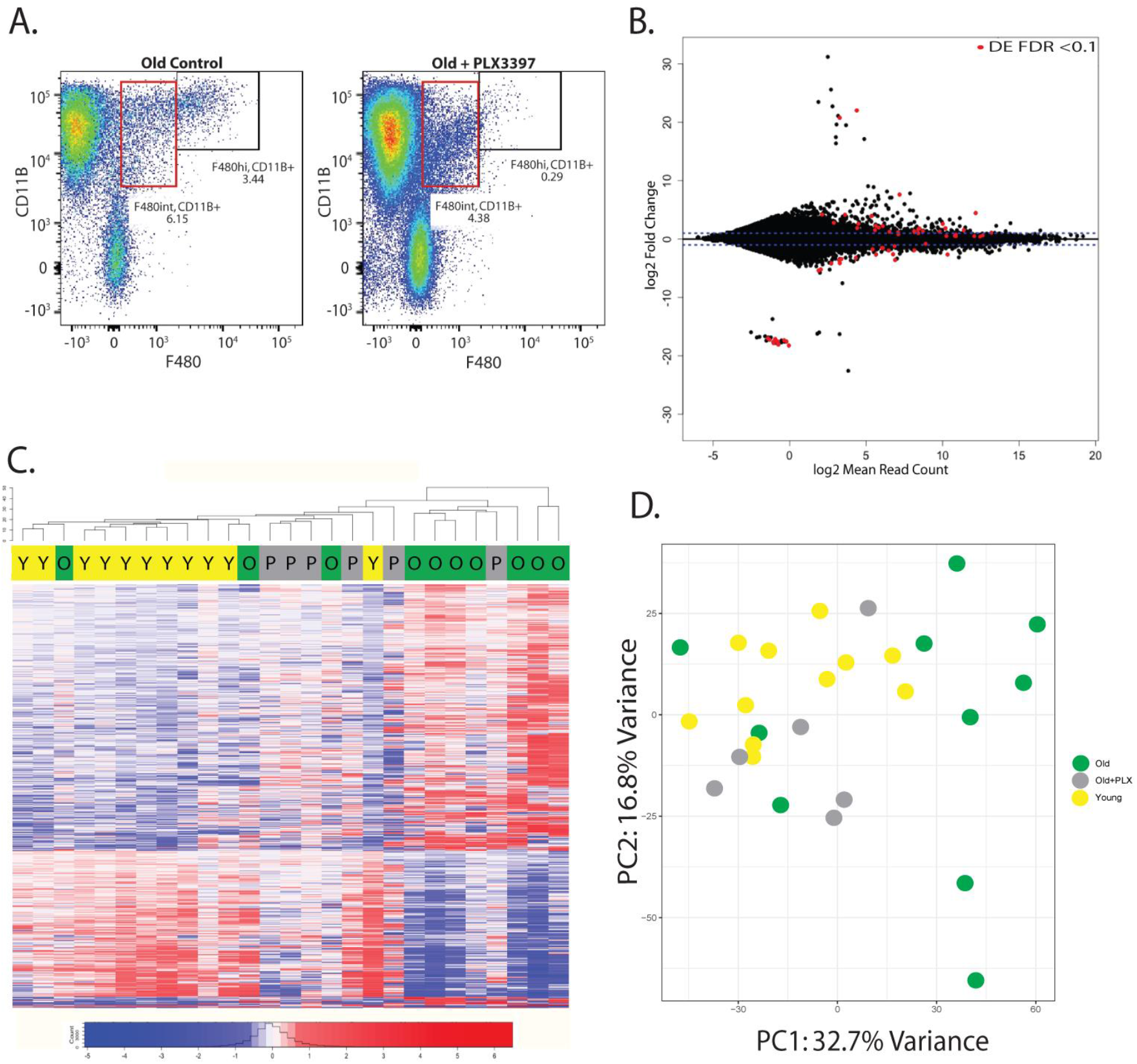
Improved fracture healing in treated old mice is associated with transcriptionally “younger” macrophages. (A) PLX3397 treatment during fracture healing resulted in significant decrease in macrophage quantity within the fracture callus, as analyzed through flow cytometry. (B) RNA-seq analysis demonstrates 64 genes were significantly differentially expressed in old macrophages treated with PLX3397 compared to young (red dots) (FDR <0.1). (C) Hierarchal clustering demonstrates old macrophages treated with PLX3397 (gray) cluster between young (yellow) and old (green) control macrophages. (D) Principal component analysis of differential gene expression in macrophages demonstrates close clustering of young (yellow) mice with old mice treated with PLX3397 (gray).

### 3.5. Improved fracture healing is associated with transcriptionally “younger” macrophages

To understand how the transcriptional profile of callus macrophages changes with PLX3397 treatment, we collected macrophages from the fracture callus of mice treated with PLX3397 at 3 days post-fracture. RNA-seq analysis demonstrated that only 64 genes were significantly differentially expressed in old macrophages from mice treated with PLX3397 compared to young macrophages; 27 genes were up-regulated and 28 genes were down-regulated more than 2-fold (Fig. 5B). Hierarchical clustering by Euclidean distance demonstrates that macrophages from old mice treated with PLX3397 cluster between macrophages from young and old mice without treatment (Fig. 5C). Principal component analysis further demonstrates that the macrophages from old mice treated with PLX3397 cluster closely with the young mice compared to the old with less transcriptomic heterogeneity (Fig. 5D). This suggests that the inflammatory macrophages that are recruited to the bone fracture are substantially different between old and young mice, but the remaining tissue resident macrophages demonstrate less age-related changes.

## 4. Discussion

The results from this study demonstrate that an aged macrophage phenotype is detrimental to fracture healing. Using an unbiased next generation sequencing approach, we demonstrate differences in the gene expression signatures of macrophages that infiltrated the fracture site in young and old mice. Macrophages from old mice have a more M1, pro-inflammatory gene signature than macrophages from young animals. Further, we demonstrated that a pharmacologic (PLX3397) leading to a decrease in macrophages recruitment to the fracture site of old mice improves fracture healing outcomes. In older mice treated with PLX3397, macrophages that are present within the callus appear transcriptionally “younger”, suggesting that the detrimental age-related changes may occur in infiltrating macrophages.

The delayed fracture healing we observed in the elderly mice compared to young adults here (Fig. 1) is similar to our previous findings [17]. Other groups have shown delayed healing with decreased callus size and decreased bone volume at multiple time points post fracture in old mice compared to young [18, 42]. Substantial alterations in inflammation may affect fracture healing in aged animals. For example, inflammation induced with Lipopolysaccharide led to decreased callus strength in young animals [10], and delayed healing in aged animals has been directly associated with inflammatory dysregulation within the callus in aged mouse models of fracture healing [24, 43]. Other studies have demonstrated an association of systemic inflammatory dysregulation, as a result of increased age or disease, with poor fracture healing outcomes in humans and animal experiments [4–6, 44]. In fact, we have shown that transplantation of juvenile bone marrow into lethally irradiated middle-aged animals stimulates bone fracture healing, and this is associated with decreased inflammation [24]. This work was subsequently confirmed [45]. However, the inflammatory cells responsible for the stimulatory effect remain largely unknown.

Recent work has suggested that macrophages may underlie the dysregulation of inflammation seen during fracture healing in aged animals. Our preliminary research demonstrated that reducing the influx of inflammatory macrophages into the callus of aged animals stimulated healing [38]. Other work has advanced this observation and supports the idea that aging macrophages are deleterious to healing [45, 24]. Interestingly, depletion of macrophages in healthy young mice has also been shown to be deleterious to fracture healing [30, 32, 33]. Collectively, this work supports the important role for macrophages in fracture healing and the deleterious effect of age-related changes to macrophages on fracture healing outcomes.

The age-related changes in fracture healing does not appear to be a function of significant differential inflammatory cell recruitment. We observed similar numbers of immune cells infiltrating the fracture callus in young and old mice (Fig. 2). T cells may contribute to fracture healing largely through the recruitment and activation of osteoclasts [46]. The aging immune system is characterized by decreased naive T cell quantity and weaker activation in elderly populations compared to young [47]. However, the quantity of T cells within the fracture callus did not differ by age (Fig. 2A). Similarly, the quantity of F4/80+ macrophages did not differ by age; however, a sub-population of macrophages, identified as F480+ Ly6C-, was increased at day 1 post fracture in young animals but this difference was not apparent by day 3 (Fig. 2C). Macrophages that are F4/80 positive and Ly6C negative have been suggested to be anti-inflammatory or M2-like macrophages [41, 48, 49], suggesting that the M2-like macrophages were present in the fracture callus of young mice to a greater extent at earlier time points than in old mice. The significance of this subtle alteration is not known.

At day 10 we observed increased B-cells in the fracture callus of young mice compared to old mice (Fig. 2A). During fracture healing B cells regulate osteoclast activity through the expression of osteoprotegerin [46], and interactions between B cells and macrophages have been demonstrated to regulate inflammation during infection [50, 51]. While these are intriguing observations the specific role of B-cells in mediating effects of age on fracture healing are unexplored, but the novelty of these cells in an inflammatory response to injury is potentially interesting.

The importance of macrophages to fracture healing and bone regeneration is beginning to emerge [33, 45, 52]. Different macrophage sub-types have been proposed to mediate separate functions during healing. Pro-inflammatory, or M1-type, macrophages appear early in fracture healing and produce pro-inflammatory cytokines (IL-1, IL-6, TNFα, iNOS) that further propagate the inflammatory response [53]. These cells likely phagocytize remnants of dead tissues to debride the fracture site. Later, a switch to anti-inflammatory, or M2-type, macrophages occurs within the fracture callus. M2 macrophages initiate down regulation of the inflammatory response with production of IL-10 and TGFβ [53, 54]. M2 macrophages also promote tissue repair through the production of growth factors (TGFβ, PDGF, VEGF) [55]. However, *in vivo* phenotyping of macrophages is complex, and given the plasticity of macrophages, the M1/M2 distinctions are likely not dichotomous and probably represent poles on a large spectrum of macrophage phenotypes. Here, RNA-seq analysis allowed an unbiased analysis of the transcriptomic differences of macrophages within the fracture callus of young and old mice. Macrophages from the fracture callus of old mice were transcriptionally distinct from macrophages of young mice (Fig. 3). Further, a M1/M2 gene expression signature, comprised of 14 selected genes for cytokines and cell markers associated with traditional M1 and M2 phenotypes, was distinct between age groups and able to differentiate macrophages from old mice versus young. Here, macrophages from old mice demonstrated expression of more pro-inflammatory or M1 genes suggesting that old macrophages contribute to the pro-inflammatory phenotype evident in elderly populations.

With the demonstrated pro-inflammatory, and potentially deleterious, phenotype of aged macrophages, we wanted to understand the effect of limiting macrophage recruitment into the fracture callus. PLX3397 treatment prevented macrophage recruitment and resulted in improved fracture healing in old mice (Fig. 4). The magnitude of change appears small in the PLX3397 treated old mice when compared to total callus volume at 10 days post fracture. However, the change represents a 35% increase in bone volume in the treated group compared to age-matched controls, which is consistent with other bone fracture research in mice that shows improvements in bone volume within a fracture callus of 25-50% using other experimental agents to improve fracture healing [56–58].

RNA-seq analysis demonstrated that 1222 genes were significantly differently expressed in macrophages in old mice compared to young (Fig. 3A), and this quantity is important considering the breadth of biological and disease processes that these genes are associated with (Fig. 3B). In old mice, when recruitment of macrophages was inhibited with PLX3397 the number of significantly differentially expressed genes between old and young macrophages were reduced by 95% (Fig. 5B), and the treated old mice clustered between the young and old controls (Fig. 5C). These findings suggest the presence of a more youthful macrophage population after PLX3397 treatment in old mice.

One potential “youthful” macrophage population is osteomacs. Osteomacs are resident tissue macrophages in bone and have been shown to co-localize with osteoblasts and contribute to osteogenesis [52, 59]. Specific depletion of osteomac populations in vivo was shown to be deleterious during both intramembranous and endochondral fracture repair processes [60]. After inhibition of M-CSF with PLX3397, the infiltrating macrophages were decreased and this is accompanied by a decrease in inflammation mediated by macrophages [38, 61]. The pharmacological effect of PLX3397 works largely on infiltrating inflammatory macrophages by antagonizing CSF1R and preventing the monocyte to macrophage differentiation. Therefore, we suspect that the resident osteomacs are less affected by PLX3397. The improved fracture healing in old mice treated with PLX3397 could be a result of decreased inflammatory macrophages and/or an expansion or activation of a more youthful and beneficial osteomac population. Further work is needed to understand the age-related changes to osteomacs and their contribution to fracture healing.

A potentially important observation that we made using RNA-seq analysis is that macrophages from old animals were much more heterogenous than those from young animals (Fig. 3C). The heatmaps demonstrated more homogenous gene expression levels between the individual young mice than old animals. The extent of heterogeneity in old mice is present despite all mice being from the same genetic background, sourced from the same laboratory and colony, housed in similar environments, and samples prepared on the same day. Complex disease processes and biological traits, including aging, are often defined by a heterogenous phenotypic presentation [62, 63, 64]. Heterogenous changes are present across many aspects of the biology of aging with an impact that is not fully understood. Age-related genetic heterogenicity could result from cumulative effects from the environment or an unknown mechanism [65]. How to properly analyze the heterogenicity present in large genetic datasets is not clearly defined. Largely, the heterogeneity is accepted as normal and sample size is increased so that differences may be detected. However, further research would be better aimed at understanding the biological cause and significance of such variation, as the increased variance in old animals may be a substantial contributor to phenotypic outcomes. Further, this may aid in identifying at risk individuals who would benefit from individualized treatment plans.

## 5. Conclusion

In summary, this study characterizes the cellular immune response during bone fracture healing and demonstrates age-related changes at the cellular level that are reflective of the altered physiology present in elderly populations. Robust transcriptional differences differentiated macrophages infiltrating the callus of old mice compared to young. The old macrophages demonstrated a pro-inflammatory M1 macrophage phenotype. The aged macrophage phenotype was detrimental to fracture healing outcomes as fracture healing was improved in old mice when aged macrophages were inhibited from accumulating in the callus during fracture healing. The therapeutic targeting of macrophages during fracture healing may be an effective therapy to improve fracture healing outcomes in elderly populations. Finally, understanding the variance and its underlying mechanism(s) may contribute significantly to directed treatments of the patient with musculoskeletal injuries

## Acknowledgements

This work was supported by the National Institutes of Health grant number R01AG046282 and the UCSF Core Center for Musculoskeletal Biology and Medicine (P30AR066262). We thank Andrea Barczak, Walter Eckalbar, and Michael Adkisson for RNA sequencing. We thank Gina Baldoza for laboratory support.

## Supplementary Figures and Tables

**Supplementary Figure 1:**
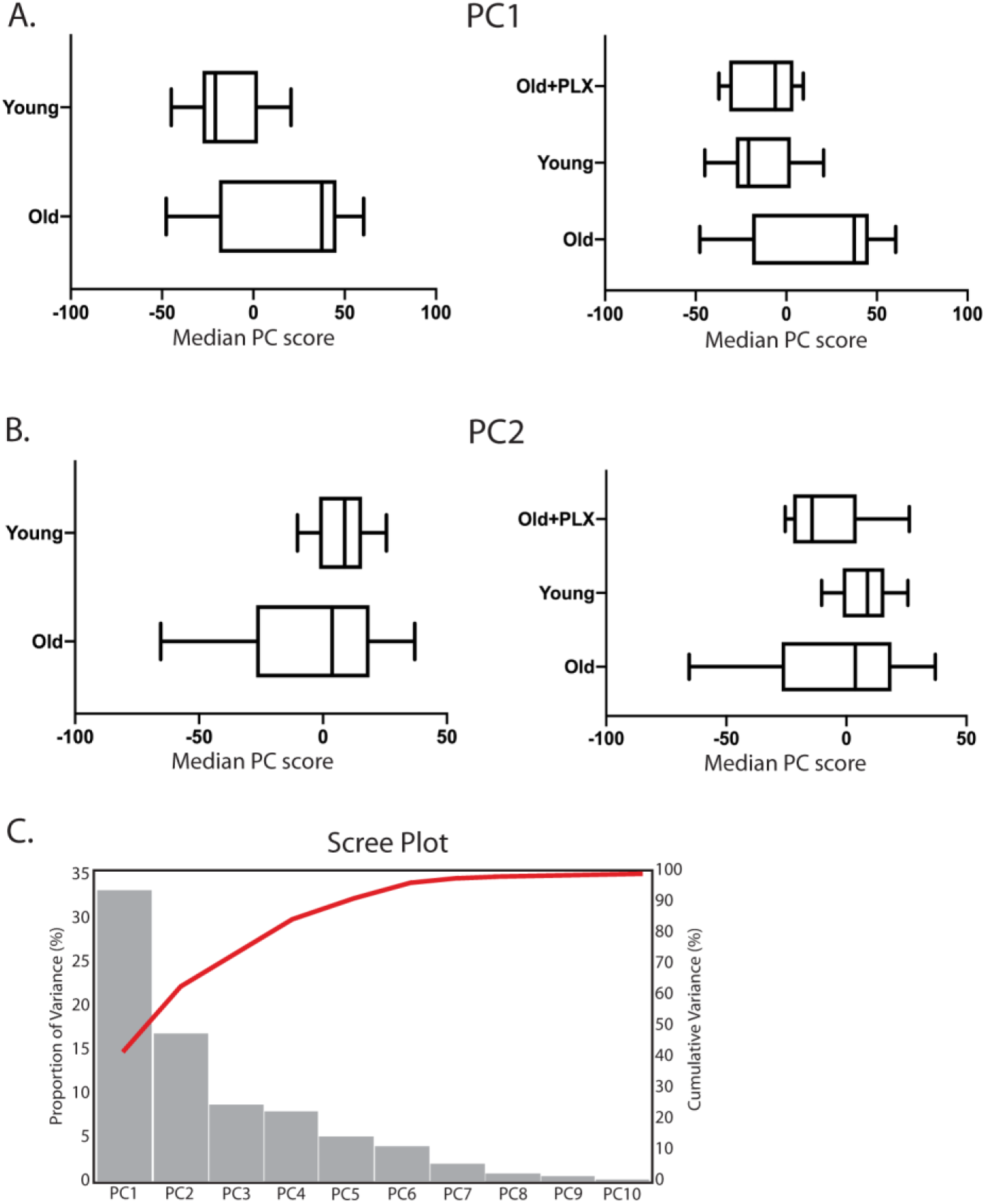
Plot of the individual PC scores along PC1 (A) and PC2 (B) of young, old, and old mice treated with PLX3397. PC1 separates old from young (A). The variance of PC scores is greater across both PC1 and PC2 in old mice compared to young. (C) Scree plot demonstrates that the majority of the variance (56.9%) is accounted for in PC1 and PC2.

**Supplementary Table 1:** Gene list of the upper and lower 2% of eigenvalues along PC1

| Upper 2% |  | Lower 2% |  |
| --- | --- | --- | --- |
| Gene | Eigenvalue | Gene | Eigenvalue |
| Igkv15-103 | 0.0584 | Gpnmb | -0.0425 |
| Ighg2c | 0.0556 | Selenop | -0.0426 |
| Ighv7-1 | 0.0547 | Slc6a8 | -0.0428 |
| Igkc | 0.0537 | P2ry6 | -0.0429 |
| Ighg2b | 0.0531 | Pdlim4 | -0.0432 |
| Igkj5 | 0.0439 | Tppp3 | -0.0436 |
| Ighv6-3 | 0.0435 | Lpl | -0.0437 |
| Igkv5-39 | 0.0430 | Fcgr1 | -0.0440 |
| Igkv1-117 | 0.0426 | Cav1 | -0.0443 |
| BC100530 | 0.0426 | Igf1 | -0.0445 |
| Stfa2l1 | 0.0425 | Pid1 | -0.0446 |
| Igkv1-88 | 0.0418 | Tmem37 | -0.0447 |
| Igkv6-32 | 0.0388 | H2-Ab1 | -0.0448 |
| Asprv1 | 0.0385 | Cx3cr1 | -0.0449 |
| Stfa2 | 0.0383 | Gdf15 | -0.0449 |
| Igkj2 | 0.0368 | Il10 | -0.0449 |
| Ighv14-2 | 0.0356 | Mmp14 | -0.0450 |
| Aspa | 0.0351 | Ednrb | -0.0451 |
| Stfa1 | 0.0343 | H2-Aa | -0.0452 |
| Ighv11-2 | 0.0343 | Ifi27l2a | -0.0453 |
| Gm5483 | 0.0335 | C3ar1 | -0.0458 |
| Pou2af1 | 0.0334 | H2-Eb1 | -0.0461 |
| Iglc2 | 0.0325 | Ccr5 | -0.0465 |
| Igha | 0.0323 | Stab1 | -0.0468 |
| Ighj4 | 0.0322 | Olfml3 | -0.0469 |
| Ighv5-16 | 0.0316 | Aif1 | -0.0477 |
| Hoxa9 | 0.0309 | Emp1 | -0.0477 |
| Ctla2a | 0.0309 | Amz1 | -0.0482 |
| Cacna1h | 0.0307 | Abcc3 | -0.0483 |
| Gm27252 | 0.0305 | Pmp22 | -0.0487 |
| Ighm | 0.0301 | Apoe | -0.0489 |
| Gimap6 | 0.0301 | Pltp | -0.0489 |
| D630045J12Rik | 0.0298 | Cd36 | -0.0494 |
| Stfa3 | 0.0297 | Spp1 | -0.0495 |
| Il5ra | 0.0296 | Ccl12 | -0.0497 |
| Hoxa7 | 0.0288 | Dab2 | -0.0497 |
| Fam110c | 0.0287 | Gas6 | -0.0497 |
| Saa3 | 0.0287 | Ifi205 | -0.0498 |
| Pdgfrb | 0.0286 | Nxpe5 | -0.0500 |
| Dmwd | 0.0284 | Lgmn | -0.0507 |
| Spns2 | 0.0284 | Ccl8 | -0.0519 |
| Mpl | 0.0279 | Pf4 | -0.0520 |
| Igkj4 | 0.0279 | Flrt3 | -0.0524 |
| Cxcr2 | 0.0278 | C1qc | -0.0526 |
| Ighj2 | 0.0277 | Folr2 | -0.0527 |
| Iglv1 | 0.0277 | Siglec1 | -0.0539 |
| Retnlg | 0.0274 | Ms4a14 | -0.0539 |
| Igkv5-43 | 0.0273 | Cxcl1 | -0.0545 |
| Il17rb | 0.0273 | Ppbp | -0.0548 |
| Ighv9-3 | 0.0270 | Sdc3 | -0.0551 |
| Scrg1 | 0.0269 | C1qb | -0.0558 |
| Ighj1 | 0.0268 | Cbr2 | -0.0570 |
| Ighv1-15 | 0.0266 | Pdpn | -0.0582 |
| Myct1 | 0.0265 | Arg1 | -0.0589 |
| Crispld2 | 0.0265 | C1qa | -0.0596 |
| Muc13 | 0.0265 | Ccl7 | -0.0598 |
| Cd55 | 0.0265 | Cd209a | -0.0598 |
| Epx | 0.0264 | Ms4a7 | -0.0610 |
| Itga2b | 0.0263 | Mrc1 | -0.0656 |
| Serpina3g | 0.0263 | Fcrls | -0.0737 |

